# STING restrains calcium-dependent microglial phagocytosis and protects against neuronal loss and cognitive impairment during epileptogenesis

**DOI:** 10.64898/2026.06.25.734646

**Authors:** Yuka Kasahara, Hideyuki Nakashima, Satoshi Miyashita, Taichi Umeyama, Yasuhiro Nakano, Soichiro Kumamoto, Kazuhiko Kawata, Keisuke Imabayashi, Yoshihiro Baba, Koji Kobiyama, Ken J Ishii, Takahiro Sato, Yoshikazu Johmura, Mikio Hoshino, Kinichi Nakashima

## Abstract

Epileptogenesis is accompanied by robust neuroinflammation, yet the molecular pathways linking innate immune activation to neuronal dysfunction remain incompletely defined. Given the established pro-inflammatory role of the stimulator of interferon genes (STING), we initially hypothesized that its loss would attenuate neuroinflammatory responses during epileptogenesis. Contrary to our expectation, we found that STING deficiency instead amplified microglial activation. Using a kainic acid mouse model of temporal lobe epilepsy (TLE), we show that STING-deficient microglia exhibit pronounced lysosomal expansion and enhanced phagocytic engulfment of neurons, leading to increased hippocampal neuronal loss and cognitive impairment. Mechanistically, STING deficiency increased the expression and altered the subcellular distribution of stromal interaction molecule 1 (STIM1), an endoplasmic reticulum Ca²⁺ sensor that mediates store-operated calcium entry (SOCE), resulting in dysregulated intracellular calcium dynamics and elevated SOCE activity. Pharmacological inhibition of SOCE reduced microglia–neuron interactions and microglial phagocytic engulfment, improved neuronal survival in the CA3 region of the hippocampus, and rescued cognitive deficits following status epilepticus. Collectively, these findings redefine STING as a negative regulator of microglial activation and identify a previously unrecognized STING–STIM1–SOCE axis that constrains calcium-dependent microglial phagocytosis during epileptogenesis, highlighting microglial calcium signaling as a potential therapeutic target in TLE.

## Introduction

Epilepsy is one of the most prevalent brain disorders, affecting an estimated 65 million people worldwide^1^. It is frequently accompanied by psychiatric and cognitive comorbidities, which substantially reduce quality of life^2,3^. Despite the availability of antiepileptic drugs, approximately one-third of patients develop drug-resistant epilepsy (DRE)^4,5^. Temporal lobe epilepsy (TLE), the most common focal epilepsy in adults, accounts for a large proportion of DRE cases and is characterized by recurrent seizures with an onset arising from limbic structures, including the hippocampus and amygdala^6^. Because a substantial proportion of patients develop DRE despite available treatments, these clinical challenges highlight the urgent need for disease-modifying therapies targeting epileptogenic mechanisms rather than symptomatic seizure suppression.

The gradual process by which normal brain tissue develops epilepsy (epileptogenesis) provides a critical window for therapeutic intervention^7,8^. Increasing evidence implicates neuroinflammation as a central driver of this process^9,10^. While acute inflammatory responses can contribute to tissue repair and homeostasis, sustained or excessive inflammation leads to maladaptive remodeling and neuronal dysfunction in epilepsy and other neurological disorders^11^. Microglia, the resident immune cells of the central nervous system, play a central role in this process by adopting activated states and releasing inflammatory mediators that amplify downstream signaling cascades^12^. However, the molecular mechanisms that regulate microglial responses during epileptogenesis remain incompletely defined.

Neuroinflammation in epilepsy is driven, in part, by activation of innate immune pattern-recognition receptors in response to pathogen- and damage-associated molecular patterns^13–16^. In line with our ongoing efforts to elucidate mechanisms underlying epileptogenesis^17^, we previously demonstrated that microglial Toll-like receptor 9 (TLR9) signaling attenuates seizure-induced aberrant neurogenesis in the adult hippocampus^18^, highlighting a modulatory role for innate immune pathways in epileptogenesis. Extending this paradigm, recent studies have identified DNA-mediated immune signaling as a key mechanism linking neuronal stress to neuroinflammatory responses in epilepsy. In particular, mitochondrial DNA (mtDNA) leakage from stressed neurons activates the cyclic GMP–AMP synthase (cGAS)–STING pathway, thereby promoting neuron–glia crosstalk and driving epileptic pathology^19^. Together, these observations position the cGAS–STING axis as a central mechanism coupling endogenous danger signals to microglial activation during epileptogenesis.

Given the established pro-inflammatory function of STING, we initially hypothesized that STING deficiency would attenuate neuroinflammatory responses following status epilepticus. To test this hypothesis, we investigated the role of STING in regulating microglial activation during epileptogenesis and its impact on neuronal survival and cognitive function using a mouse model of TLE.

## Results

### STING restrains microglial accumulation and lysosomal expansion during epileptogenesis

Previous studies have reported increased STING expression in microglia in human epileptic brains and experimental models of epilepsy^20,21^; however, its functional role in epileptogenesis remains incompletely understood. Consistent with previous reports, STING expression was elevated in microglia within the CA1 and CA3 regions of the mouse hippocampus following KA-induced SE (Extended Data Fig. 1a–c). To define the role of STING in epileptogenesis, we performed bulk RNA sequencing (RNA-seq) of hippocampi from wild-type (WT) and STING-KO mice with or without SE. STING deficiency resulted in a distinct transcriptional profile following SE (Fig. 1a). Genes associated with immune responses were markedly upregulated, whereas genes involved in synaptic function and cognition were downregulated, indicating enhanced neuroinflammatory activation accompanied by impaired neuronal function. Unsupervised clustering revealed clear segregation of STING-deficient mice after SE, particularly at 7 days, indicating progressive divergence in transcriptional responses (Fig. 1b). Differential expression analysis identified 3121 genes significantly altered between WT and STING-KO mice at this time point (Fig. 1c), the majority of which were associated with innate immune pathways and microglial function (Fig. 1d and Extended Data Fig. 2a-g). Notably, IFN-I transcripts were undetectable in both genotypes, regardless of SE, whereas several downstream components of canonical STING signaling were paradoxically upregulated in STING-deficient mice at 7 days after SE (Extended Data Fig. 2h-l), suggesting a noncanonical regulatory role for STING in this context.

**Fig. 1.**
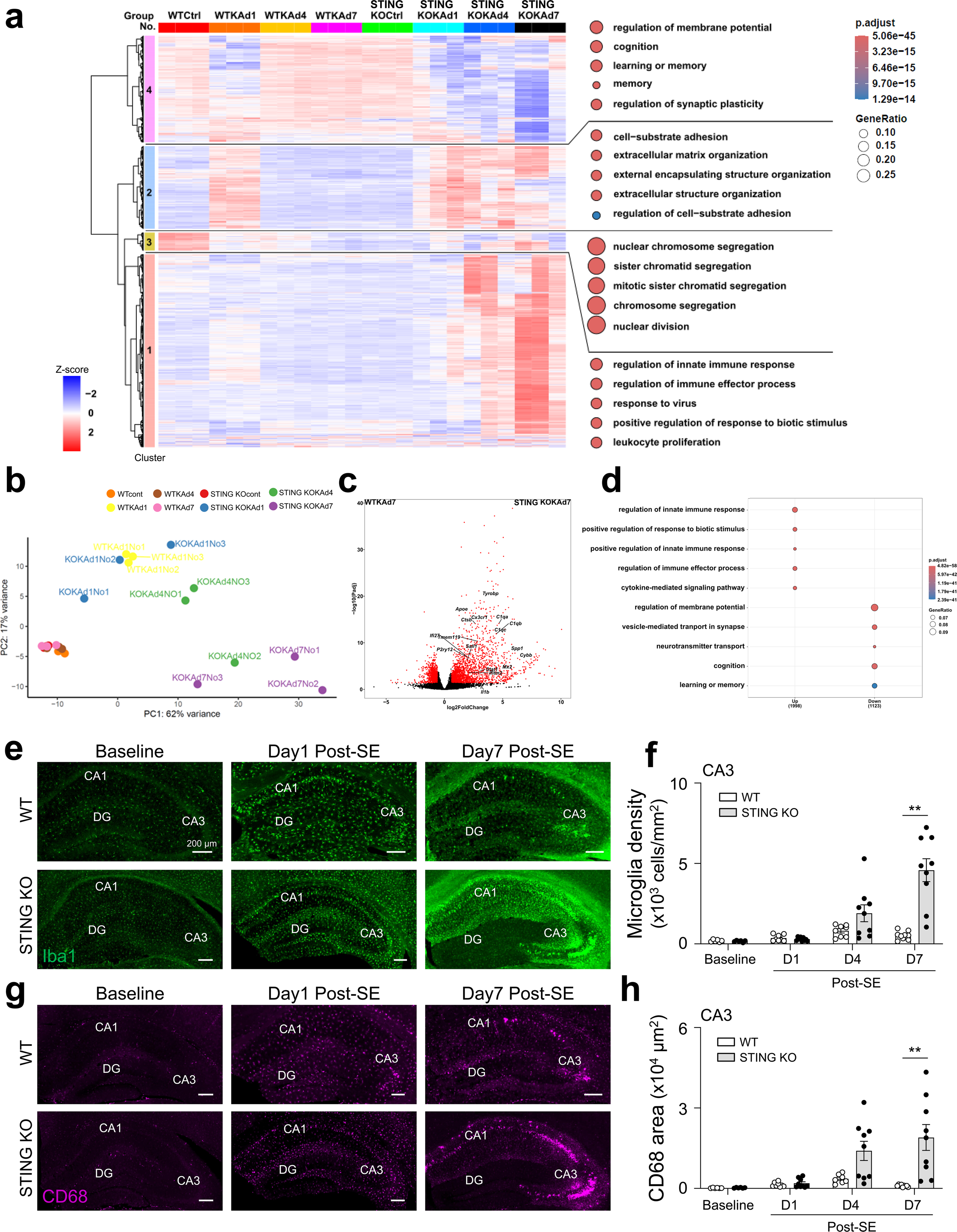
STING deficiency enhances microglial accumulation and lysosomal expansion during epileptogenesis. **a**, Heat map of gene expression and dot plots showing GO terms associated with each cluster. Dot plots display selected enriched GO terms for each cluster. Colors in the heat map represent normalized, scaled expression values. Dot color indicates adjusted P values obtained from a hypergeometric over-representation test with Benjamini–Hochberg correction for multiple comparisons. **b**, Principal component analysis (PCA) of hippocampal transcriptomes from WT and STING-KO mice. **c**, Volcano plot showing DEGs between WT and STING-KO mice at 7 days post-SE. **d**, Dot plots showing selected enriched GO terms associated with DEGs, including 1998 upregulated and 1123 downregulated genes. Dot color indicates adjusted P values following Benjamini–Hochberg correction, and dot size corresponds to gene ratio. **e**, **g**, Representative images of the hippocampus from WT and STING-KO mice under naïve baseline conditions and at 1 and 7 days post-SE. Iba1 (green, e) and CD68 (magenta, g). **f**, **h**, Microglial density (f) and CD68-positive area (h) in the CA3 region. Data are presented as mean ± s.e.m. Statistical analysis was performed using two-way ANOVA followed by Tukey’s multiple-comparison test (**P < 0.01). *n* = 3 mice (a–d) and 6–9 regions from 3 mice per group (f, h).

In light of these transcriptional changes, we next assessed microglial responses in the hippocampal pyramidal layer. B oth WT and STING-deficient mice exhibited increased Iba1-positive microglial density and CD68-positive lysosomal area following SE, which were more pronounced in STING-deficient mice (Fig. 1e–h and Extended Data Fig. 1d–g). At 7 days after SE, STING-KO mice exhibited approximately 4- and 9-fold increases in microglial density, and 9- and 20-fold increases in CD68-positive area in the CA1 and CA3 regions, respectively, compared to WT mice (Fig. 1f, h and Extended Data Fig. 1h, i). In contrast, microglial density and CD68 expression were comparable between genotypes under basal conditions. Together, these data demonstrate that STING restrains microglial accumulation and lysosomal activation during early epileptogenesis.

### S TING restrains microglial engulfment of neurons during epileptogenesis

Pronounced phagolysosomal expansion was observed in the pyramidal cell layer of the CA3 region, suggesting enhanced microglial phagocytic activity. To determine whether microglia actively engage in neuronal phagocytosis, we quantified interactions between CD68-positive lysosomal structures and NeuN-positive neurons at 7 days after SE. These interactions were classified into four stages, ranging from no contact to full engulfment with reference to previous studies^20^ (Extended Data Fig. 3a). In WT mice, most neurons exhibited no or minimal interaction with CD68-positive signals. In contrast, STING-KO mice displayed markedly increased phagocytic engagement, with approximately 61 % of neurons displaying moderate CD68 coverage and 12 % undergoing complete engulfment (Fig. 2a, b). Consistent with these findings, pronounced neuronal loss was observed in regions enriched with CD68-high microglia in STING-deficient mice, whereas WT mice showed minimal neuronal loss. (Fig. 2c, d and Extended Data Fig. 3b). Neuronal densities on the CA regions were comparable between genotypes under naïve condition (Extended Data Fig. 3c-e), indicating that neuronal loss is specifically induced following SE in the absence of STING.

**Fig. 2.**
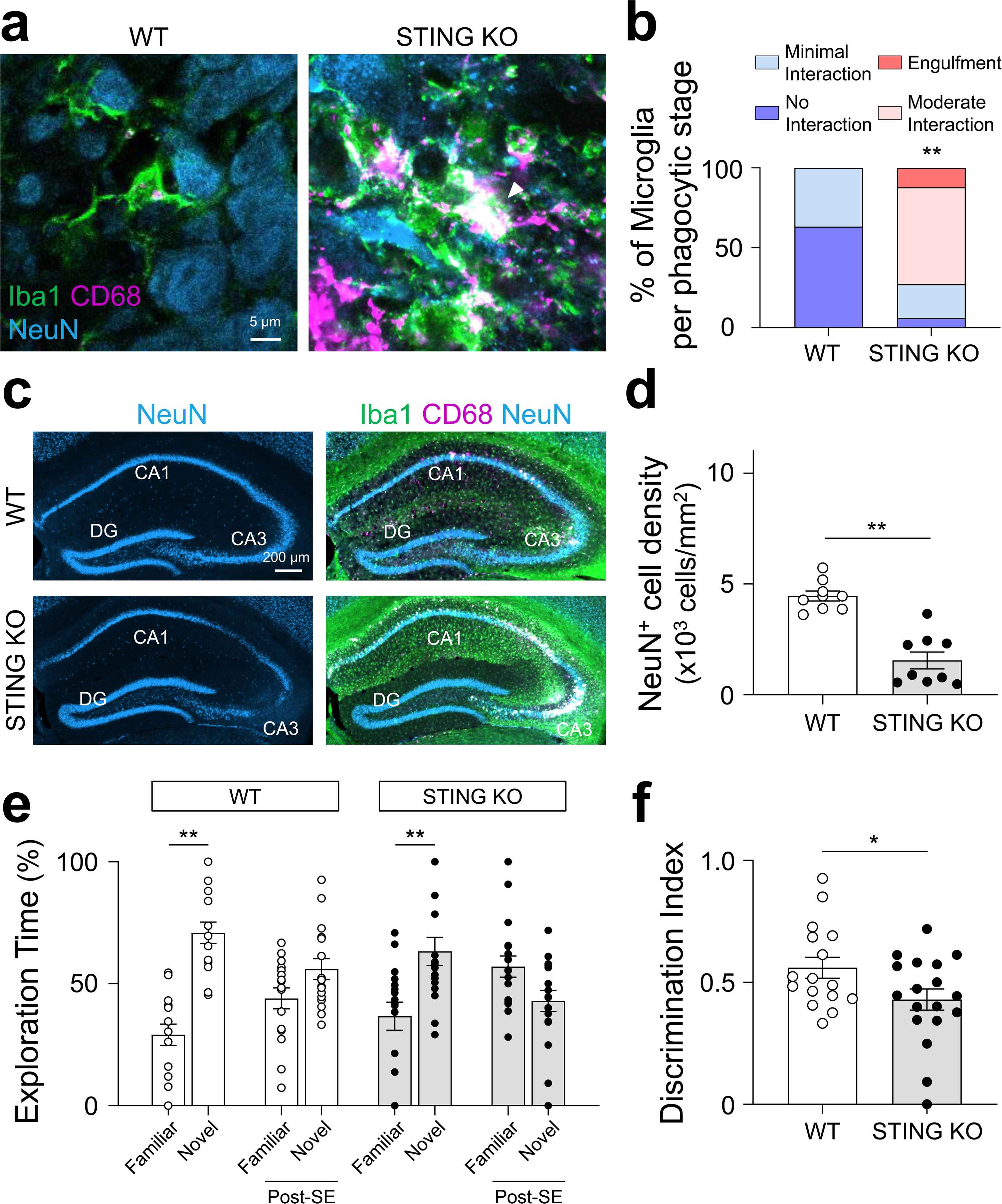
STING deficiency promotes microglial phagocytosis of neurons and exacerbates SE-induced cognitive impairment. **a**, Representative images of microglia in the CA3 pyramidal cell layer of WT and STING-KO mice at 7 days post-SE. Iba1 (green), CD68 (magenta), and NeuN (blue). White arrowheads indicate NeuN-positive signals overlapping with CD68-positive structures. **b**, Distribution of neurons across phagocytic stages. **c**, Representative images of the hippocampus from WT and STING-KO mice at 7 days post-SE. Iba1 (green), CD68 (magenta), and NeuN (blue). **d**, NeuN⁺ cell density in the CA3 region at 7 days post-SE. **e**, Percentage of time spent exploring familiar and novel objects in WT and STING-KO mice under naïve conditions and at 2 weeks post-SE. **f**, Discrimination index in WT and STING-KO mice at 2 weeks post-SE. Data are presented as mean ± s.e.m. Statistical analysis was performed using one-way ANOVA followed by Tukey’s multiple-comparison test (e) or an unpaired Student’s t-test (f). **P < 0.01 and *P < 0.05. *n* = 30–33 cells from 3 mice (b), 6–9 regions from 3 mice per group (d), and 16–18 mice (e, f).

To determine whether these pathological changes affect cognitive function, we performed the novel object recognition (NOR) task, which depends on hippocampal function, including CA3-dependent processes^22,23^. During the familiarization phase, no preference for identical objects was observed in any group (Extended Data Fig. 3f, g). In the subsequent test phase, control mice preferentially explored the novel object. In contrast, SE-exposed mice failed to show a significant preference (Fig. 2e). Notably, STING-deficient mice exhibited a reduced discrimination index compared to WT mice following SE (Fig. 2f). Total distance traveled during the test phase was comparable between genotypes (Extended Data Fig. 3h), indicating that impaired performance was not attributable to altered locomotor activity.

Together, these findings demonstrate that STING limits microglia-mediated neuronal engulfment and protects against seizure-induced neuronal loss and cognitive impairment during early epileptogenesis.

### Microglia-specific STING deletion recapitulates enhanced microglial engulfment of neurons and associated deficits following SE

To investigate whether the observed phenotypes are mediated by microglial STING, we generated microglia-specific STING conditional knockout mice using *Hexb^CreERT2^* mice^24^ crossed with *STING^fl/fl^*mice. Tamoxifen (TAM) administration at postnatal days P7 and P9 resulted in efficient and sustained deletion of STING in microglia into adulthood (Extended Data Fig. 4a-c). At 7 days after SE, microglia-specific STING-deficient (STING cKO) mice exhibited markedly increased microglial accumulation and lysosomal expansion in the CA3 region compared to *Hexb^CreERT2^* (control) mice (Fig. 3a–d). Microglial density and CD68-positive area were increased by 15-fold and 41-fold, respectively, indicating robust activation of microglial phagolysosomal pathways. Consistent with these observations, microglial engulfment of neurons was substantially increased in STING cKO mice. Quantitative analysis revealed that 55% of neurons exhibited moderate CD68 coverage and 32% were fully engulfed (Fig. 3e, f), highlighting markedly elevated phagocytic activity compared to controls. The increased microglial engulfment was accompanied by pronounced neuronal loss in regions enriched with CD68-positive microglia, whereas control mice showed minimal neuronal loss (Fig. 3g, h). Notably, these phenotypes were more pronounced than those observed in global STING-deficient mice. This may reflect known characteristics of the *Hexb^CreERT2^* model, including subtle alterations in microglial gene expression^24^, or compensatory STING signaling in non-microglial cell types. Nevertheless, these findings indicate that microglial STING restrains microglia-mediated neuronal engulfment and protects against neuronal loss during epileptogenesis.

**Fig. 3.**
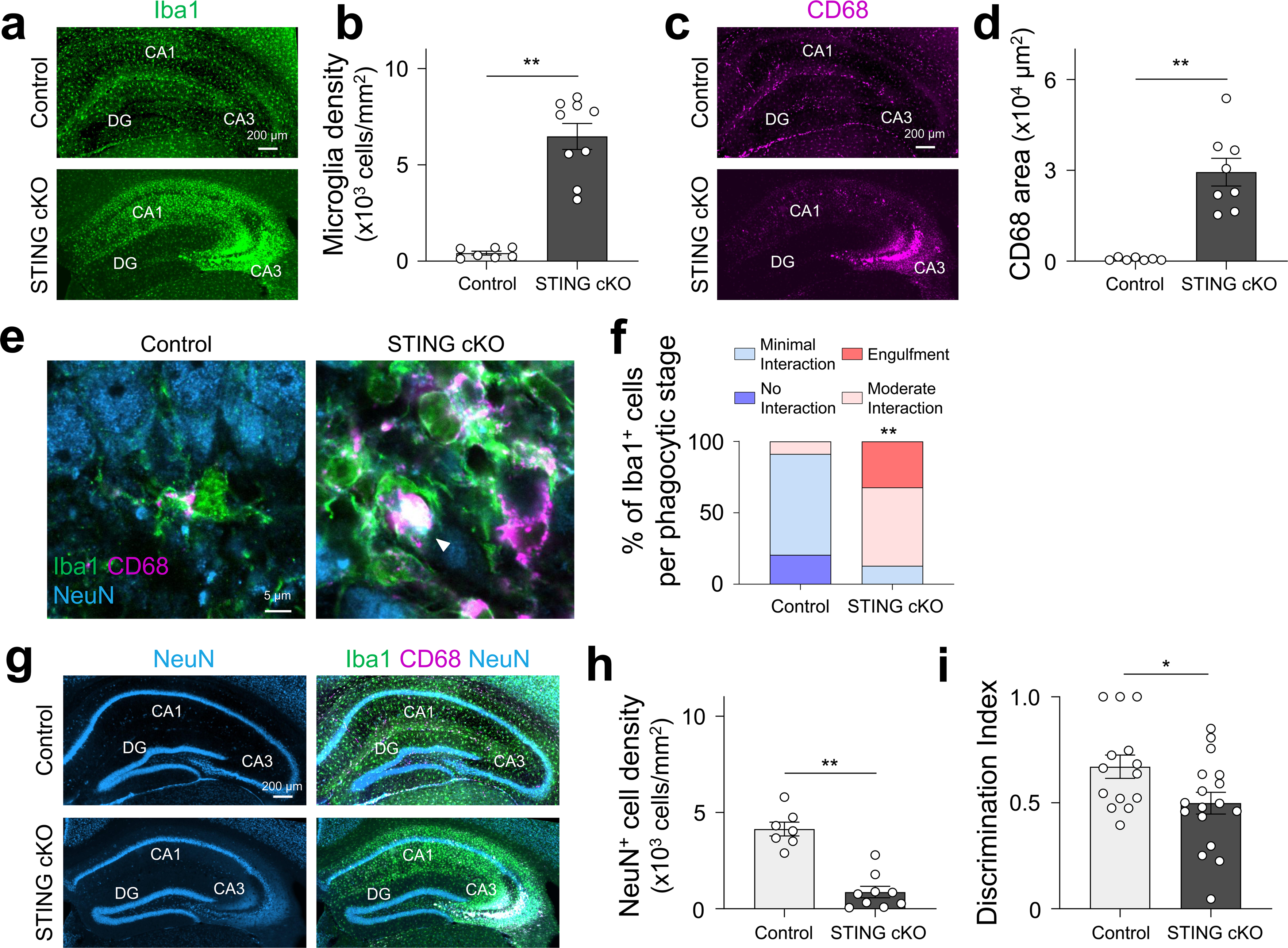
Microglial STING deficiency drives microglial hyperactivation and impairs cognition during epileptogenesis. **a**, **c**, Representative images of the hippocampus from control and STING cKO mice at 7 days post-SE. Iba1 (green, a) and CD68 (magenta, c). **b**, **d**, Microglial cell density (b) and CD68 area (d) in the CA3 region. **e**, Representative images of microglia in the CA3 pyramidal cell layer of control and STING cKO mice at 7 days post-SE. Iba1 (green), CD68 (magenta), and NeuN (blue). **f**, Distribution of neurons across phagocytic stages. White arrowheads indicate NeuN-positive signals overlapping with CD68-positive structures. **g**, Representative hippocampal images from control and STING cKO mice at 7 days post-SE. Iba1 (green), CD68 (magenta), and NeuN (blue). **h**, NeuN⁺ cell density in the CA3 region. **i**, Discrimination index in NOR task at 2 weeks post-SE. Data are presented as mean ± s.e.m. Statistical analysis was performed using an unpaired Student’s t-test. **P < 0.01. *n* = 6–9 regions from 3 mice per group (b, d, h), 31–34 cells from 3 mice (f), and 14–18 mice (i).

Consistent with these histological changes, STING cKO mice exhibited impaired cognitive performance in the NOR task. Compared to control mice, STING cKO mice displayed a reduced discrimination index (Fig. 3i). The discrimination index exceeded chance level (0.5; one-sample t-test) in control mice but not in STING cKO mice. Locomotor activity was comparable between genotypes (Extended Data Fig. 4d), indicating that the observed deficits were not attributable to altered movement.

Together, these findings demonstrate that microglial STING is a critical regulator of microglial phagocytic activity and is required to limit neuronal loss and cognitive impairment during epileptogenesis.

### STING deficiency enhances STIM1–Orai1 signaling during epileptogenesis and drives aberrant calcium influx in microglia

To elucidate the mechanism by which STING regulates microglial function during epileptogenesis, we next investigated STING activation dynamics in microglia following SE. In the resting state, STING directly interacts with the Ca²⁺ sensor, STIM1, retaining STING in an inactive state at the ER membrane^25^. Upon activation by cytosolic DNA, STING translocates to the ER–Golgi intermediate compartment (ERGIC), where it recruits TANK-binding kinase 1 (TBK1) and interferon regulatory factor 3 (IRF3)^26,27^. Following SE, the co-localized volume of STING with the cis-Golgi marker GM130 was increased at 1 day but decreased at 7 days compared with naïve WT mice, whereas STING–STIM1 co-localization showed the opposite pattern (Fig. 4a–d). These results indicate dynamic STING trafficking, with early activation and ERGIC translocation followed by reassociation with STIM1 at later stages of epileptogenesis.

**Fig. 4.**
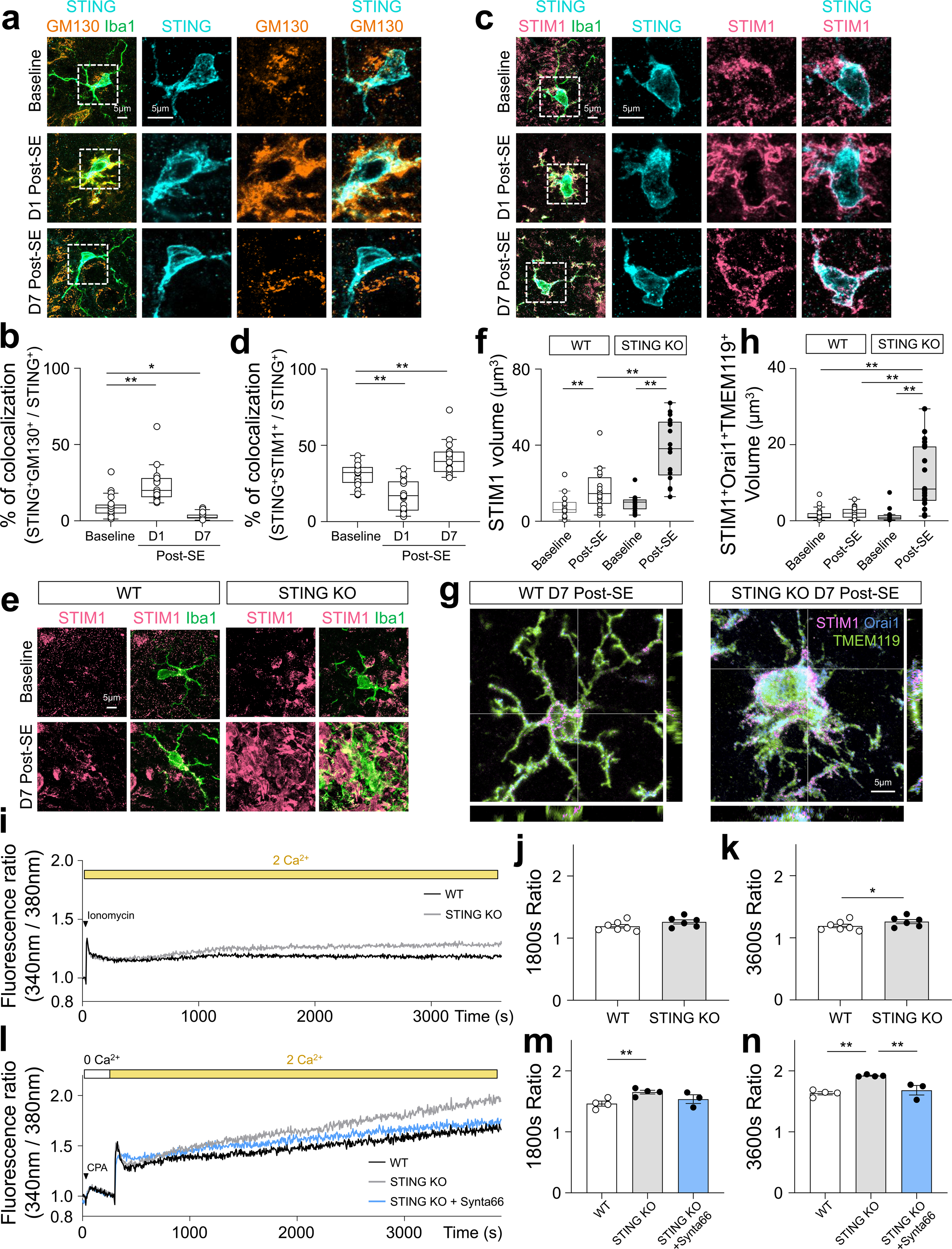
STING constrains STIM1–Orai1–dependent calcium signaling in microglia during epileptogenesis. **a**, Representative images of microglia from WT mice under naïve baseline conditions and at 1 and 7 days post-SE. Iba1 (green), STING (cyan), and GM130 (orange). **b**, Percentage of STING^+^GM130^+^ volume relative to total STING^+^ volume in individual microglia within the CA3 region. **c**, Representative images of the microglia from WT mice under naïve baseline conditions and at 1 and 7 days post-SE. Iba1 (green), STING (cyan), and STIM1 (pink). **d**, Percentage of STING^+^STIM1^+^ volume relative to total STING^+^ volume in individual microglia. **e**, Representative images of microglia from WT and STING-KO mice under naïve conditions and at 7 days post-SE. Iba1 (green) and STIM1 (pink). **f**, Quantification of STIM1 signal volume in individual microglia. **g**, Three-dimensional reconstructed images of microglia from WT and STING-KO at 7 days post-SE. TMEM119 (green), Orai1 (blue), and STIM1 (pink). **h**, Colocalized volume of STIM1 and Orai1 within TMEM119-positive microglia **i**, Time course of ionomycin-induced Ca²⁺ responses, expressed as the 340/380 nm fluorescence ratio in WT and STING-KO primary cultured microglia. **j**, **k**, Fluorescence ratio at 1800 s (j) and 3600 s (k) after recording onset. **l**, Time course of CPA-evoked SOCE in WT and STING-KO primary cultured microglia with or without Synta66 treatment. **m**, **n**, Fluorescence ratio at 1800 s (m) and 3600 s (n) after recording onset. For box plots, center lines indicate medians, box limits indicate the first and third quartiles, and whiskers extend to 1.5× the interquartile range according to the Tukey method. Statistical analysis was performed using one-way ANOVA with Tukey’s multiple-comparison test (b, d, f, h, m, n) or an unpaired Student’s t-test (j, k). **P < 0.01 and *P < 0.05. *n* = 20—21 microglia from 3—4 mice (b), 19—20 microglia from 3 mice (d), 17—20 microglia from 3 mice (f), 20—22 microglia from 3—5 mice (h), 7 independent cultures (i—k), and 3—4 independent cultures (l—n).

We next examined whether STING regulates STIM1 expression. STIM1 expression was increased in microglia following SE in WT mice, consistent with the previous report of increased STIM1 expression in hippocampal tissue from patients with mTLE^28^, and the expression was further elevated in STING-KO mice (Fig. 4e, f). STIM1 redistributes to ER–plasma membrane junctions upon ER Ca²⁺ depletion, where it activates Orai1 to induce store-operated calcium entry (SOCE)^29,30^. Consistent with this mechanism, Orai1 expression was increased in STING-KO mice, and co-localization of STIM1 with Orai1 and the microglial membrane marker TMEM119 was enhanced following SE (Fig. 4g, h and Extended Data Fig. 5a, b), indicating potentiation of STIM1–Orai1 signaling.

To determine the functional consequences of these changes, we measured intracellular Ca²⁺ dynamics in primary microglial cultures. Ionomycin-induced Ca²⁺ responses showed comparable peak amplitudes between genotypes but exhibited sustained elevation in STING-deficient microglia over time (Fig. 4i–k). We next directly assessed SOCE activity by depleting ER Ca²⁺ stores using cyclopiazonic acid (CPA) under Ca²⁺-free conditions, followed by Ca²⁺ readdition. While the initial Ca²⁺ influx was similar between genotypes, the sustained phase of Ca²⁺ entry was significantly enhanced in STING-KO microglia (Fig. 4l-n). Similar results were observed in UDP-stimulation (Extended Data Fig. 5c-f). Importantly, this enhanced SOCE response was suppressed by the Orai channel inhibitor, 3-fluoro-pyridine-4-carboxylic acid (2,5-dimethoxy-biphenyl-4-yl)-amide (Synta66)^31^ (Fig. 4n), confirming the involvement of Orai-dependent mechanisms.

To further support this involvement, we re-analyzed publicly available transcriptomic data from Orai1-deficient microglia^32^. Genes downregulated in Orai1-deficient microglia upon treatment with thapsigargin and phorbol 12,13-dibutyrate, which induce SOCE and activate Ca^2+^-dependent transcriptional programs, substantially overlapped with genes upregulated in STING-KO mice following SE, whereas minimal overlap was observed with genes upregulated in WT mice (Extended Data Fig. 5g–i). These findings suggest that STING deficiency induces transcriptional programs that are inversely related to Orai1-dependent Ca²⁺ signaling. Together, these results demonstrate that STING constrains STIM1–Orai1–dependent SOCE and likely contributes to the maintenance of intracellular Ca²⁺ homeostasis in microglia during epileptogenesis.

### Pharmacological inhibition of SOCE rescues neuronal loss and cognitive impairment during epileptogenesis

To evaluate the therapeutic potential of targeting calcium-dependent microglial activation, we next investigated whether inhibition of SOCE ameliorates SE-induced phenotypes in STING-KO mice in vivo. Given the limited validation of Synta66 for central delivery, we used BTP2 (3,5-bis(trifluoromethyl)pyrazole-2; YM-58483), a potent SOCE inhibitor with established central activity, and administered it orally to model a clinically relevant route of administration^33,34^. In STING-KO mice, BTP2 treatment significantly reduced microglial accumulation and lysosomal expansion in the CA3 pyramidal cell layer at 7 days after SE (Fig. 5a–d).

**Fig. 5.**
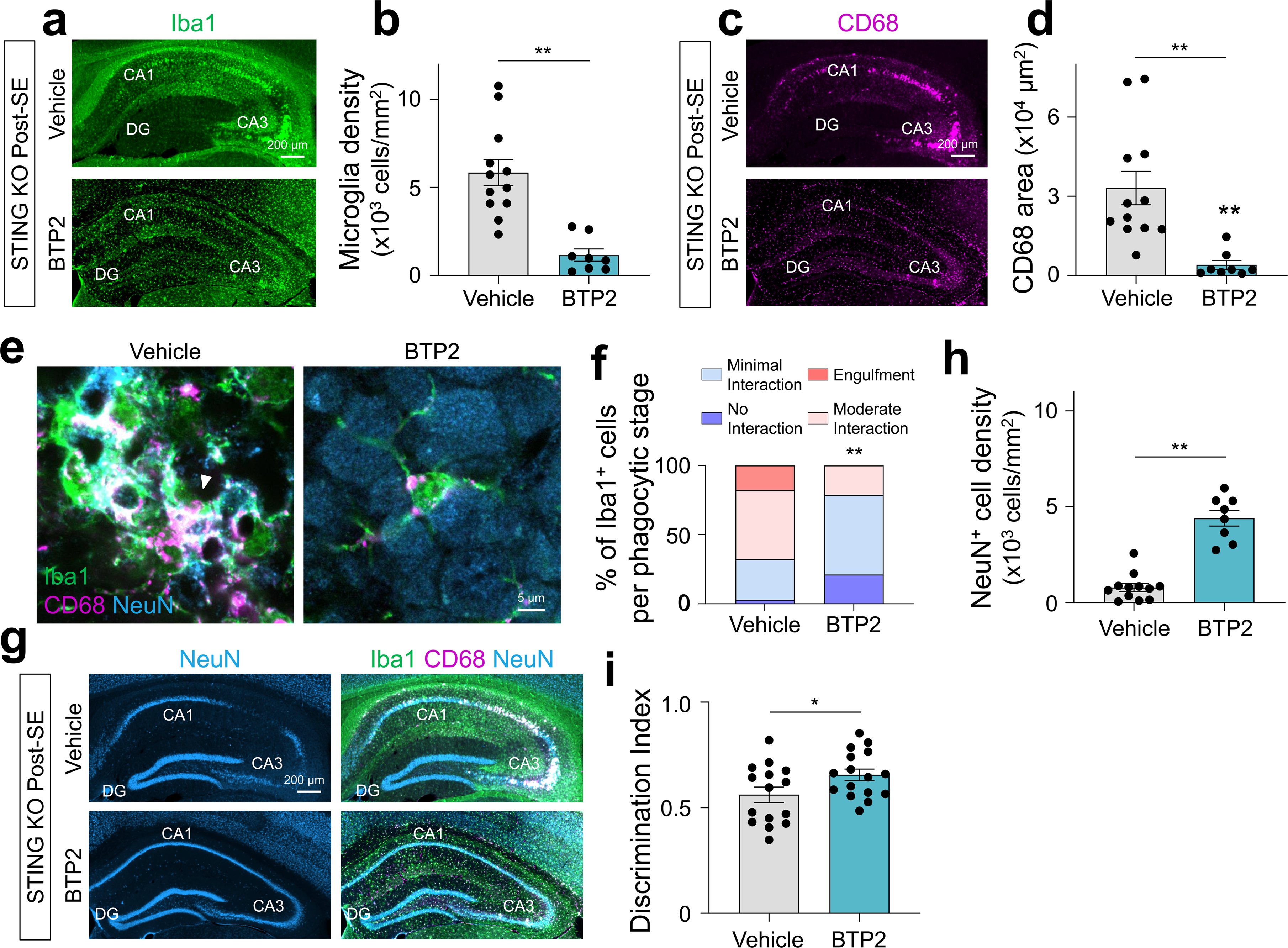
Pharmacological inhibition of SOCE rescues neuronal loss and cognitive impairment during epileptogenesis. **a, c**, Representative images of the hippocampus from vehicle- or BTP2-treated STING-KO mice at 7 days post-SE. Iba1 (green, a) and CD68 (magenta, c). **b**, Microglial density in the CA3 region. **d**, CD68-positive area in the CA3 region. **e**, Representative images of microglia in the CA3 pyramidal cell layer from vehicle- or BTP2-treated STING-KO mice at 7 days post-SE. Iba1 (green), CD68 (magenta), and NeuN (blue). White arrowheads indicate NeuN-positive signals overlapping with CD68-positive structures. **f**, Distribution of neurons across phagocytic stages. **g**, Representative hippocampal images from vehicle- or BTP2-treated STING-KO mice at 7 days post-SE. Iba1 (green), CD68 (magenta), and NeuN (blue). **h**, NeuN⁺ cell density in the CA3 region. **i**, Discrimination index in the NOR task at 2 weeks post-SE. Data are presented as mean ± s.e.m. Statistical analysis was performed using an unpaired Student’s t-test (b, d, h, i). **P < 0.01 and *P < 0.05. *n* = 8–12 regions from 3 mice (b, d, h), 33–34 cells from 3 mice (f), and 15–16 mice (i).

Consistent with these effects, microglia–neuron interactions were markedly attenuated. In vehicle-treated STING-KO mice, the majority of neurons exhibited moderate to full CD68 coverage, with 18% of neurons fully engulfed. In contrast, BTP2 treatment markedly shifted this distribution, increasing the proportion of neurons with no or minimal CD68 contact and eliminating fully engulfed neurons (Fig. 5e, f), indicating reduced microglial phagocytic activity. This reduction in microglial phagocytosis was accompanied by preservation of neuronal integrity, as evidenced by preserved NeuN-positive cell density in the CA3 region following SE (Fig. 5g, h).

We next assessed whether these histological improvements translate into functional recovery. In the NOR task, BTP2-treated STING-KO mice showed a significantly increased discrimination index that exceeded chance level (0.5; one-sample t-test), whereas vehicle-treated STING-KO mice failed to show object preference (Fig. 5i). Locomotor activity was unaffected (Extended Data Fig. 6a), indicating that improved performance reflected cognitive recovery.

Overall, these findings demonstrate that pharmacological inhibition of SOCE rescues microglia-mediated neuronal loss and cognitive impairment, identifying microglial calcium signaling as a potential therapeutic target during epileptogenesis.

## Discussion

The cGAS–STING signaling pathway is widely recognized as a central driver of inflammatory responses across diverse pathological conditions^35^. In the context of epilepsy, recent studies have shown that inhibition of cGAS–STING signaling attenuates neuroinflammation and reduces seizure phenotypes in models driven by genetic or mitochondrial dysfunction^19,21^. In contrast to this prevailing view, our study identifies an unexpected and opposing role for STING in epileptogenesis. We show that STING deficiency exacerbates neuroinflammation, enhances microglia-mediated neuronal engulfment, and worsens cognitive impairment following status epilepticus. These findings redefine STING as a negative regulator of microglial activation during early epileptogenesis.

This apparent discrepancy highlights the context-dependent functions of STING. While STING is classically regarded as a pro-inflammatory signaling adaptor, accumulating evidence indicates that it can also contribute to immune homeostasis and tissue protection^36,37^. Consistent with this concept, our results demonstrate that STING restrains excessive microglial activation despite ongoing inflammatory stimuli. Differences in disease etiology, temporal dynamics, and upstream signaling pathways, including the relative contribution of the cGAS axis and mitochondrial stress signals, may underlie the divergent roles of STING observed across models of epilepsy and neuroinflammation.

Our data further reveal that STING undergoes dynamic subcellular trafficking during epileptogenesis. We observe transient translocation of STING to the ERGIC in the acute phase, followed by reassociation with the ER-resident Ca²⁺ sensor STIM1 at later stages. This temporal pattern suggests that STING activity is tightly regulated and may transition from an initial signaling phase to a homeostatic regulatory state. The interaction between STING and STIM1 therefore emerges as a potentially critical checkpoint linking innate immune activation to intracellular Ca²⁺ signaling.

At the cellular level, STING deficiency induces a hyperactivated microglial state characterized by increased lysosomal expansion and pronounced phagocytic engulfment of neurons. Notably, transcriptional analyses revealed the coexistence of activation-associated and homeostatic gene expression programs, including P2ry12, Tmem119, and Cx3cr1. This profile deviates from canonical disease-associated microglia (DAM) signatures^38^, which are typically defined by loss of homeostatic gene expression together with induction of inflammatory and phagocytic programs. These findings suggest that microglial activation during early epileptogenesis represents a distinct, dynamic state and that STING plays a key role in constraining this transition. Future single-cell transcriptomic analyses will be required to determine whether these transcriptional changes reflect a discrete microglial state or shifts in the relative abundance of existing microglial populations.

Although our study primarily focused on microglia-mediated phagocytosis, additional mechanisms may contribute to neuronal degeneration following SE. STING deficiency markedly increased expression of pro-inflammatory genes (Fig, 1c, d), which may promote neuronal dysfunction both directly and through induction of neurotoxic astrocytes^9,39^. In addition, STING signaling in non-microglial cell types, including peripheral immune cells, may influence disease progression^40,41^. While microglia-specific STING deletion recapitulated the observed phenotypes, we cannot fully exclude contributions from other cell types, as *Hexb^CreERT2^*-mediated recombination is not entirely restricted to microglia^42^. Future studies using more refined cell-type-specific approaches are warranted to dissect these contributions further.

Intracellular Ca²⁺ signaling is a key regulator of microglial activation during epileptogenesis^20^; however, the mechanisms governing its dysregulation remain incompletely defined. Here, we identify STING as an upstream regulator of microglial Ca²⁺ homeostasis. STING deficiency enhanced STIM1 and Orai1 expression, promoted their spatial association, and led to a sustained elevation of intracellular Ca²⁺ via augmented SOCE. This was supported by functional analyses showing increased CPA-and UDP-evoked SOCE responses in STING-deficient microglia, which were abolished by the Orai inhibitor Synta66. A previous study has shown that STIM1 interacts with STING to retain it in the ER, thereby linking innate immune signaling to Ca²⁺ homeostasis^25^. Our findings extend this concept by demonstrating that STING constrains STIM1 redistribution and STIM1–Orai1–dependent Ca²⁺ influx. These results establish STING as an upstream regulator of a Ca²⁺-dependent signaling pathways that govern microglial activation and phagocytic function during epileptogenesis. A limitation of the present study is that functional evidence was primarily obtained through pharmacological inhibition of SOCE. Future studies using microglia-specific genetic manipulation of STIM1 in microglia will be required to define the precise mechanisms underlying STING-dependent Ca²⁺ regulation.

Importantly, targeting the microglial STING–STIM1–Orai1 axis may have translational relevance. Several CRAC/Orai inhibitors, including CM4620 (Auxora, zegocractin), have already advanced into clinical trials^43,44^, demonstrating the feasibility of therapeutically modulating SOCE signaling. However, because STIM1 and Orai1 are broadly expressed, application to neurological disorders will require cell–type–specific strategies. Notably, previous studies have shown that neuronal SOCE plays essential roles in synaptic plasticity and seizure susceptibility, and that its disruption exacerbates seizure phenotypes^45,46^. These observations indicate that SOCE signaling exerts distinct functions across cell types, with neuronal SOCE supporting circuit stability while excessive microglial SOCE promoting maladaptive inflammatory responses.

In summary, our findings redefine STING as a homeostatic regulator of microglial activation and identify a previously unrecognized STING–STIM1–Orai1 axis that constrains Ca²⁺-dependent phagocytic activity during epileptogenesis. Altogether, these findings identify microglial calcium signaling as a central therapeutic target and suggest that selective modulation of the STING–SOCE axis represents a potential disease-modifying strategy for temporal lobe epilepsy.

## Materials and Methods

### Mice

All efforts were made to minimize animal suffering and reduce the number of animals used. Mice were housed under a 12-h light/12-h dark cycle with ad libitum access to food and water. All experiments were conducted in accordance with the guidelines for animal experimentation of Kyushu University and complied with the National Institutes of Health Guide for the Care and Use of Laboratory Animals. WT C57BL/6 mice and *STING^fl/fl^* mice were obtained from Japan SLC and The Jackson Laboratory, respectively. STING-KO mice were generated from *Tmem173^tm1Camb^* (KOMP) Mbp ES cell line (JM8A3.N1) obtained from Knockout Mouse Project (KOMP) Repository (California, U.S.A.)^47^. *Hexb^CreERT2^* mice^24^ were generously provided by Dr. Takahiro Masuda (Kyushu University). *Hexb^CreERT2^* mice were crossed with *STING^fl/fl^*mice to generate STING cKO mice. To induce Cre recombination, tamoxifen (Sigma-Aldrich) was administered at postnatal day 7 (P7) and P9 (0.4 mg TAM dissolved in 20 μl corn oil per injection), as the previous study^24^. BTP2 (Selleck, S8380) was suspended in saline containing 1% DMSO and 1% Cremophor EL (Nacalai Tesque, 09727-14) and orally administered at 10 mg/kg every other day until the experimental endpoint.

### Induction of status epilepticus with kainic acid

To minimize variability in seizure susceptibility and mortality associated with sex differences following KA administration^48^, only male mice were used throughout the study. SE was induced in 8-week-old mice by intraperitoneal administration of kainic acid (KA; Tocris, 7065). WT and STING-KO mice received KA at 30mg/kg and, control and STING cKO mice received 20mg/kg. Behavioral seizures were assessed according to the Racine scale^49^. To reduce variability in seizure severity and neuropathological outcomes, only mice exhibiting continuous seizures reaching Racine stage 3 or higher were included in subsequent analyses. Because convulsive SE is characterized by generalized clonic seizures^50^, only mice displaying clear clonic seizures were used for further experiments. Mice that did not exhibit clonic seizures within the first hour after the initial KA injection received additional intraperitoneal injections of KA (5 mg/kg) every 20 min until the first clonic seizure was observed.

### mRNA isolation and cDNA library preparation for bulk RNA-seq

mRNA was isolated from dissected hippocampi of WT and STING-KO mice using the Dynabeads mRNA DIRECT Micro Purification Kit (Invitrogen, 61021). Purified mRNA was used for cDNA library preparation with the NEBNext Ultra Directional RNA Library Prep Kit for Illumina (New England Biolabs, E7420) according to the manufacturer’s instructions. Briefly, polyadenylated mRNA was isolated using Dynabeads Oligo(dT)25 (Invitrogen, 61002). mRNA was fragmented in NEBNext First Strand Synthesis Reaction Buffer by heating at 94 °C for 15 min. First-strand cDNA was synthesized from fragmented mRNA, followed by second-strand cDNA synthesis in which dUTP was substituted for dTTP. The resulting cDNA was end-repaired, dA-tailed, and ligated to NEBNext adaptors. Second-strand cDNA containing dUTP was digested using USER enzyme, and sequencing indices and barcodes were introduced by 12–15 cycles of PCR amplification. Libraries were purified using AMPure XP beads (Beckman Coulter, A63881), and library quality was assessed using a Bioanalyzer High Sensitivity DNA Kit (Agilent Technologies, 5067-4626).

### Immunocytochemistry

Mice were deeply anesthetized with isoflurane and transcardially perfused with ice-cold phosphate-buffered saline (PBS), followed by 4% paraformaldehyde (PFA). Brains were isolated, post-fixed in 4% PFA overnight at 4 °C, cryoprotected sequentially in 10% and 30% sucrose solutions for 48 h each at 4 °C, embedded in optimal cutting temperature compound (Tissue-Tek, Sakura Finetek, 25608-930), and stored at −80 °C until use. Frozen brains were coronally sectioned at 40 μm thickness using a cryostat (Leica). Immunostaining was performed as previously described. Briefly, sections were blocked for 1 h at room temperature in PBS containing 5% fetal bovine serum (FBS) and 0.3% Triton X-100, followed by incubation with primary antibodies overnight at 4 °C. After three washes with PBS (10 min each), sections were incubated with secondary antibodies for 1 h at room temperature. Nuclei were counterstained with Hoechst 33258 (Nacalai Tesque). The following primary antibodies were used: rabbit anti-Iba1 (1:500; Wako, 019-19741), goat anti-Iba1 (1:500; Abcam, ab5076), rat anti-CD68 (1:500; Bio-Rad, MCA1957), mouse anti-NeuN (1:500; clone A60, Millipore, MAB377), rabbit anti-STING (1:500; Proteintech, 19851-1-AP), mouse anti-STIM1 (1:500; clone 5A2, Novus, H00006786-M01), rabbit anti-STIM1 (1:200; clone D88E10, Cell Signaling Technology, 5668), mouse anti-Orai1 (1:200; clone 3F6H5, Novus, NBP1-75522), mouse anti-GM130 (1:200; BD Biosciences, 610822), and guinea pig anti-TMEM119 (1:500; Synaptic Systems, 400004). Secondary antibodies included CF488A donkey anti-rabbit IgG (1:500; Biotium, 20015), CF555 donkey anti-rabbit IgG (1:500; Biotium, 20038), CF488 donkey anti-mouse IgG (1:500; Biotium, 20014), CF647 donkey anti-mouse IgG (1:500; Biotium, 20046), CF568 donkey anti-rat IgG (1:500; Biotium, 20092), CF647 donkey anti-goat IgG (1:500; Biotium, 20048), and Alexa Fluor 647 goat anti-guinea pig IgG (1:500; Invitrogen, A21450). Fluorescence images were acquired using confocal laser-scanning microscopes (LSM700 and LSM800, Zeiss) equipped with a 20×/0.8 NA objective or a 63×/1.4 NA oil-immersion objective, or using a BZ-X700 fluorescence microscope (Keyence) equipped with a 10×/0.45 NA objective.

### Cell count and CD68 area analysis

To quantify microglial density, neuronal density, and CD68-positive area in the hippocampal CA1 and CA3 regions, sections were tile-scanned using a Zeiss confocal microscope with a 20× objective (1024 × 1024 pixels). A single optical plane was analyzed in ImageJ using regions of interest (ROIs) encompassing one-third of the pyramidal cell layer in the CA1 or CA3 region. ROIs were selected based on regions exhibiting the highest microglial accumulation within each section. The numbers of Iba1-positive and NeuN-positive cells within each ROI were manually counted. For CD68 quantification, uniform thresholds were applied to Iba1 and CD68 channels, and the CD68-positive area overlapping with Iba1-positive signals within the ROI was measured in ImageJ. Microglia–neuron interactions in the CA3 region were quantified as follows: (1) uniform thresholds were applied to CD68 and NeuN channels; (2) ROIs outlining individual NeuN-positive cells were generated; (3) NeuN ROIs were overlaid onto thresholded CD68 images, and the CD68-positive area within each NeuN ROI was quantified and normalized to the NeuN ROI area; and (4) overlap was classified as no interaction, minimal overlap (<10%), moderate overlap (10–80%), or engulfment (>80%).

### Signal volume analysis

To quantify STING, STIM1, GM130, Orai1, and TMEM119 signal volumes, sequential z-stack images were acquired from the CA3 region using a Zeiss confocal microscope with a 63× objective (1024 × 1024 pixels, 0.33 μm z-step interval). Image analysis was performed in ImageJ as follows: (1) a uniform threshold was applied to the Iba1 channel; (2) ROIs outlining individual Iba1-positive microglia were generated; (3) uniform thresholds were applied to STING, GM130, STIM1, and Orai1 channels, and only signals overlapping with Iba1-positive areas were analyzed; and (4) the signal volumes within each Iba1 ROI were quantified. Colocalization volumes of STING–GM130 and STING–STIM1 were normalized to STING volume. For analysis of STIM1, Orai1, and TMEM119 colocalization, uniform thresholds were applied to each channel, and colocalized volumes were quantified in ImageJ. Three-dimensional reconstruction images were performed using Imaris software.

### Novel object recognition task

Mice were handled by the experimenter and habituated to an empty arena for 10 min per day for 6–7 days before behavioral testing. The arena, constructed from opaque plastic (50 × 50 × 30 cm), was enclosed within a sound-attenuating chamber providing controlled temperature, humidity, and constant illumination. Mouse behavior was monitored and recorded using an overhead camera system. Before each task, mice were transferred to the testing room and habituated for at least 1 h. During the training phase, mice were allowed to freely explore the arena with two identical objects for 5 min. Mice were then returned to their home cages for 30 min before the test phase. During the test phase, one familiar object was replaced with a novel object, and mice were allowed to explore the arena for 5 min. Familiar and novel objects were randomly selected from three object sets for which mice showed no innate preference during preliminary testing. The arena and objects were thoroughly cleaned with 70% ethanol between trials. A discrimination index was calculated as the time spent exploring the novel object divided by the total exploration time for both objects during the test phase. Exploration behavior was defined as the mouse entering a circular zone (10 cm diameter) centered on an object. Mice that failed to explore both objects during the training phase or exclusively climbed one object during the test phase were excluded from analysis.

### Primary microglia culture

Primary microglia cultures were prepared as previously described^62^. To obtain mixed glial cells, cortex and hippocampus were dissected from P1-2 WT and STING-KO mice. The tissues were digested with papain (22.5 U/ml; Sigma, P3125) and 0.01% DNase I (SIGMA-Aldrich, D4527) at 37 °C for 20 min, followed by centrifugation at 200 × g for 5 min. Cells were resuspended in maintenance medium consisting of DMEM/Ham’s F-12 (Nacalai Tesque) supplemented with 0.1% gentamicin, 1 mM sodium pyruvate, MEM non-essential amino acids, 20% fetal bovine serum (FBS), and GM-CSF (2.5 ng/ml; R&D Systems), and plated onto T75 culture flasks (BD Falcon). The flask was placed in a 37 °C, 5% CO2 incubator and the medium was replaced the day after plating and every 3 days thereafter. After 10–14 days in vitro, microglia were detached by shaking the flasks and plated onto 96-well black-walled, clear-bottom plates coated with poly-L-lysine at a density of 4.0 - 5.0 × 10^4^ cells per well. After 30 min, the medium was replaced with maintenance medium containing 10% FBS. Primary microglia were used for experiments the following day.

### Measurement of cytosolic Ca^2+^ concentration

Cultured microglia were washed with standard extracellular solution containing 155 mM NaCl, 1 mM MgCl₂, 4.5 mM KCl, 5.6 mM D-glucose, 5 mM HEPES, 2 mM CaCl₂, and 0.025% bovine serum albumin (pH 7.4). Cells were loaded with Fura-2 AM (3 μM; Invitrogen, F1221) supplemented with Pluronic F-127 (Invitrogen, P6866) for 45 min at 37 °C in 5% CO₂. Fluorescence signals were measured at 510 nm emission with alternating excitation wavelengths of 340 and 380 nm using a FlexStation 3 microplate reader (Molecular Devices). The 340/380 nm fluorescence ratio was normalized to the mean value of the first six measurement points after recording onset. Intracellular Ca²⁺ responses were measured following application of ionomycin calcium salt (Sigma-Aldrich, I0634), cyclopiazonic acid (CPA; Sigma-Aldrich, C1530), UDP disodium salt (Selleck, S3368), and extracellular Ca²⁺ readdition. To induce store-operated calcium entry (SOCE), cells were incubated in Ca²⁺-free extracellular solution containing 0.5 mM EGTA for 5 min before recording. Synta66 (10 μM; Sigma-Aldrich, SML1949) was added beginning at the Fura-2 loading step and maintained throughout the experiment.

### Quantification and statistical analysis

Statistical analyses were performed using GraphPad Prism software. Data distribution normality and variance homogeneity were assessed before selecting statistical tests. Two-group comparisons at a single time point were analyzed using unpaired two-tailed Student’s t-tests. For comparisons involving two independent variables, two-way ANOVA followed by Tukey’s post hoc test was used. Comparisons among more than three groups at a single time point were analyzed using one-way ANOVA followed by Tukey’s post hoc test. Multiple-comparison corrections were applied where appropriate. Sample sizes were determined based on previous studies using similar experimental approaches^20^. Behavioral analyses were performed blinded to genotype, whereas image and fluorescence analyses were conducted using automated or semi-automated analysis pipelines to minimize experimenter bias.

## Acknowledgments

We appreciate the technical assistance from The Research Support Center, Research Center for Human Disease Modeling, Kyushu University Graduate School of Medical Sciences, which is partially supported by the Mitsuaki Shiraishi Fund for Basic Medical Research. Technical support for the use of the FlexStation 3 microplate reader was generously provided by Dr. Yuki Shiimura (Kurume University). This work was supported by JSPS KAKENHI (Grant Numbers JP21K15272 to Y.K.; JP23H00391 and JP25H01319 to K.N.) and by AMED (Grant Number JP20gm1310008 to K.N.).

## Author Contributions

Y.K., H.N. and K.N. conceived the project and planned the experiments. Y.K., H.N., S.M., Y.B., Y.J., M.H. and K.N. designed experiments. Y.K., H.N., S.M., T.U., Y.N. and S.K. performed experiments or analyses. Y.K., K.K. and K.I. developed experimental protocols. K.K. and K.J.I. established STING-KO mice and T.S. provided experimental resources and technical support (FlexStation 3 microplate reader). Y.K. and K.N. wrote the manuscript. All authors read and approved the manuscript.

## Notes

### Competing Interest Statement

The authors have declared no competing interest.

## References

1. Devinsky, O., et al. Epilepsy. Nat Rev Dis Primers 4, 18024 (2018).

2. Celiker Uslu, S., Yuksel, B., Tekin, B., Sariahmetoglu, H. & Atakli, D. Cognitive impairment and drug responsiveness in mesial temporal lobe epilepsy. Epilepsy Behav 90, 162–167 (2019).

3. Tellez-Zenteno, J.F., Patten, S.B., Jette, N., Williams, J. & Wiebe, S. Psychiatric comorbidity in epilepsy: a population-based analysis. Epilepsia 48, 2336–2344 (2007).

4. Chen, Z., Brodie, M.J., Liew, D. & Kwan, P. Treatment Outcomes in Patients With Newly Diagnosed Epilepsy Treated With Established and New Antiepileptic Drugs: A 30-Year Longitudinal Cohort Study. JAMA Neurol 75, 279–286 (2018).

5. Kalilani, L., Sun, X., Pelgrims, B., Noack-Rink, M. & Villanueva, V. The epidemiology of drug-resistant epilepsy: A systematic review and meta-analysis. Epilepsia 59, 2179–2193 (2018).

6. Marathe, K., et al. Resective, Ablative and Radiosurgical Interventions for Drug Resistant Mesial Temporal Lobe Epilepsy: A Systematic Review and Meta-Analysis of Outcomes. Front Neurol 12, 777845 (2021).

7. Klein, P., Kaminski, R.M., Koepp, M. & Loscher, W. New epilepsy therapies in development. Nat Rev Drug Discov 23, 682–708 (2024).

8. Ravizza, T., et al. mTOR and neuroinflammation in epilepsy: implications for disease progression and treatment. Nat Rev Neurosci 25, 334–350 (2024).

9. Vezzani, A., Balosso, S. & Ravizza, T. Neuroinflammatory pathways as treatment targets and biomarkers in epilepsy. Nat Rev Neurol 15, 459–472 (2019).

10. Li, W., Wu, J., Zeng, Y. & Zheng, W. Neuroinflammation in epileptogenesis: from pathophysiology to therapeutic strategies. Front Immunol 14, 1269241 (2023).

11. Xanthos, D.N. & Sandkuhler, J. Neurogenic neuroinflammation: inflammatory CNS reactions in response to neuronal activity. Nat Rev Neurosci 15, 43–53 (2014).

12. Benson, M.J., Manzanero, S. & Borges, K. Complex alterations in microglial M1/M2 markers during the development of epilepsy in two mouse models. Epilepsia 56, 895–905 (2015).

13. Maroso, M., et al. Toll-like receptor 4 and high-mobility group box-1 are involved in ictogenesis and can be targeted to reduce seizures. Nat Med 16, 413–419 (2010).

14. Iori, V., et al. Receptor for Advanced Glycation Endproducts is upregulated in temporal lobe epilepsy and contributes to experimental seizures. Neurobiol Dis 58, 102–114 (2013).

15. Gross, A., et al. Toll-like receptor 3 deficiency decreases epileptogenesis in a pilocarpine model of SE-induced epilepsy in mice. Epilepsia 58, 586–596 (2017).

16. Li, D. & Wu, M. Pattern recognition receptors in health and diseases. Signal Transduct Target Ther 6, 291 (2021).

17. Kasahara, Y., Nakashima, H. & Nakashima, K. Seizure-induced hilar ectopic granule cells in the adult dentate gyrus. Front Neurosci 17, 1150283 (2023).

18. Matsuda, T., et al. TLR9 signalling in microglia attenuates seizure-induced aberrant neurogenesis in the adult hippocampus. Nat Commun 6, 6514 (2015).

19. Jiang, J., et al. mtDNA leakage promotes neuron-glia crosstalk to induce epilepsy by cGAS-STING-driven neuroinflammation and serine metabolic reprogramming. Proc Natl Acad Sci U S A 123, e2522313123 (2026).

20. Umpierre, A.D., et al. Microglial P2Y(6) calcium signaling promotes phagocytosis and shapes neuroimmune responses in epileptogenesis. Neuron 112, 1959–1977 e1910 (2024).

21. Huang, Y., et al. cGAS-mediated IFN-I signaling contributes to disease progression in drug-refractory epilepsy. bioRxiv (2026).

22. Antunes, M. & Biala, G. The novel object recognition memory: neurobiology, test procedure, and its modifications. Cogn Process 13, 93–110 (2012).

23. Rolls, E.T. A quantitative theory of the functions of the hippocampal CA3 network in memory. Front Cell Neurosci 7, 98 (2013).

24. Masuda, T., et al. Novel Hexb-based tools for studying microglia in the CNS. Nat Immunol 21, 802–815 (2020).

25. Srikanth, S., et al. The Ca(2+) sensor STIM1 regulates the type I interferon response by retaining the signaling adaptor STING at the endoplasmic reticulum. Nat Immunol 20, 152–162 (2019).

26. Barber, G.N. STING: infection, inflammation and cancer. Nat Rev Immunol 15, 760–770 (2015).

27. Li, T. & Chen, Z.J. The cGAS-cGAMP-STING pathway connects DNA damage to inflammation, senescence, and cancer. J Exp Med 215, 1287–1299 (2018).

28. Steinbeck, J.A., et al. Store-operated calcium entry modulates neuronal network activity in a model of chronic epilepsy. Exp Neurol 232, 185–194 (2011).

29. Zhang, S.L., et al. STIM1 is a Ca2+ sensor that activates CRAC channels and migrates from the Ca2+ store to the plasma membrane. Nature 437, 902–905 (2005).

30. Prakriya, M. & Lewis, R.S. Store-Operated Calcium Channels. Physiol Rev 95, 1383–1436 (2015).

31. Jairaman, A. & Prakriya, M. Molecular pharmacology of store-operated CRAC channels. Channels (Austin) 7, 402–414 (2013).

32. DeMeulenaere, K.E., et al. Microglial reactivity and neuroinflammation-driven changes in motivational behaviors are regulated by Orai1 calcium channels. Sci Signal 19, eady8398 (2026).

33. Stegner, D., et al. Loss of Orai2-Mediated Capacitative Ca(2+) Entry Is Neuroprotective in Acute Ischemic Stroke. Stroke 50, 3238–3245 (2019).

34. Xiao, W., Xiao, F., Zhang, Y. & Zeng, L. BTP2, a store-operated calcium channel inhibitor, attenuates morphine antinociceptive tolerance in rats. Front Neurosci 20, 1758352 (2026).

35. Decout, A., Katz, J.D., Venkatraman, S. & Ablasser, A. The cGAS-STING pathway as a therapeutic target in inflammatory diseases. Nat Rev Immunol 21, 548–569 (2021).

36. Morch, M.T., et al. STING Signaling Deficiency Exacerbates Demyelination and Immune Infiltration in Focal EAE Lesions. NeuroSci 6(2025).

37. Sulka, K.B., et al. Microglial STING is a central safeguard against neurological decline with age. Cell Rep 44, 115749 (2025).

38. Keren-Shaul, H., et al. A Unique Microglia Type Associated with Restricting Development of Alzheimer’s Disease. Cell 169, 1276–1290 e1217 (2017).

39. Liddelow, S.A., et al. Neurotoxic reactive astrocytes are induced by activated microglia. Nature 541, 481–487 (2017).

40. Varvel, N.H., et al. Infiltrating monocytes promote brain inflammation and exacerbate neuronal damage after status epilepticus. Proc Natl Acad Sci U S A 113, E5665–5674 (2016).

41. Xu, D., et al. Peripherally derived T regulatory and gammadelta T cells have opposing roles in the pathogenesis of intractable pediatric epilepsy. J Exp Med 215, 1169–1186 (2018).

42. Frosch, M., et al. Microglia-neuron crosstalk through Hex-GM2-MGL2 maintains brain homeostasis. Nature 646, 913–924 (2025).

43. Sutton, R., et al. Zegocractin for acute pancreatitis with systemic inflammatory response syndrome: a randomized, controlled, dose-ranging, phase 2b trial. EClinicalMedicine 93, 103757 (2026).

44. Bruen, C., et al. Auxora vs. placebo for the treatment of patients with severe COVID-19 pneumonia: a randomized-controlled clinical trial. Crit Care 26, 101 (2022).

45. Hori, K., Tsujikawa, S., Novakovic, M.M., Yamashita, M. & Prakriya, M. Regulation of chemoconvulsant-induced seizures by store-operated Orai1 channels. J Physiol 598, 5391–5409 (2020).

46. Maneshi, M.M., et al. Orai1 Channels Are Essential for Amplification of Glutamate-Evoked Ca(2+) Signals in Dendritic Spines to Regulate Working and Associative Memory. Cell Rep 33, 108464 (2020).

47. Hayashi, T., et al. DAMP-Inducing Adjuvant and PAMP Adjuvants Parallelly Enhance Protective Type-2 and Type-1 Immune Responses to Influenza Split Vaccination. Front Immunol 9, 2619 (2018).

48. Li, F. & Liu, L. Comparison of kainate-induced seizures, cognitive impairment and hippocampal damage in male and female mice. Life Sci 232, 116621 (2019).

49. Racine, R.J. Modification of seizure activity by electrical stimulation. II. Motor seizure. Electroencephalogr Clin Neurophysiol 32, 281–294 (1972).

50. Wylie, T., Sandhu, D.S. & Murr, N.I. Status Epilepticus. in StatPearls (Treasure Island (FL), 2026).

