## Supplementary Figures and Legends for "STING restrains calcium-dependent microglial phagocytosis and protects against neuronal loss and cognitive impairment during epileptogenesis"

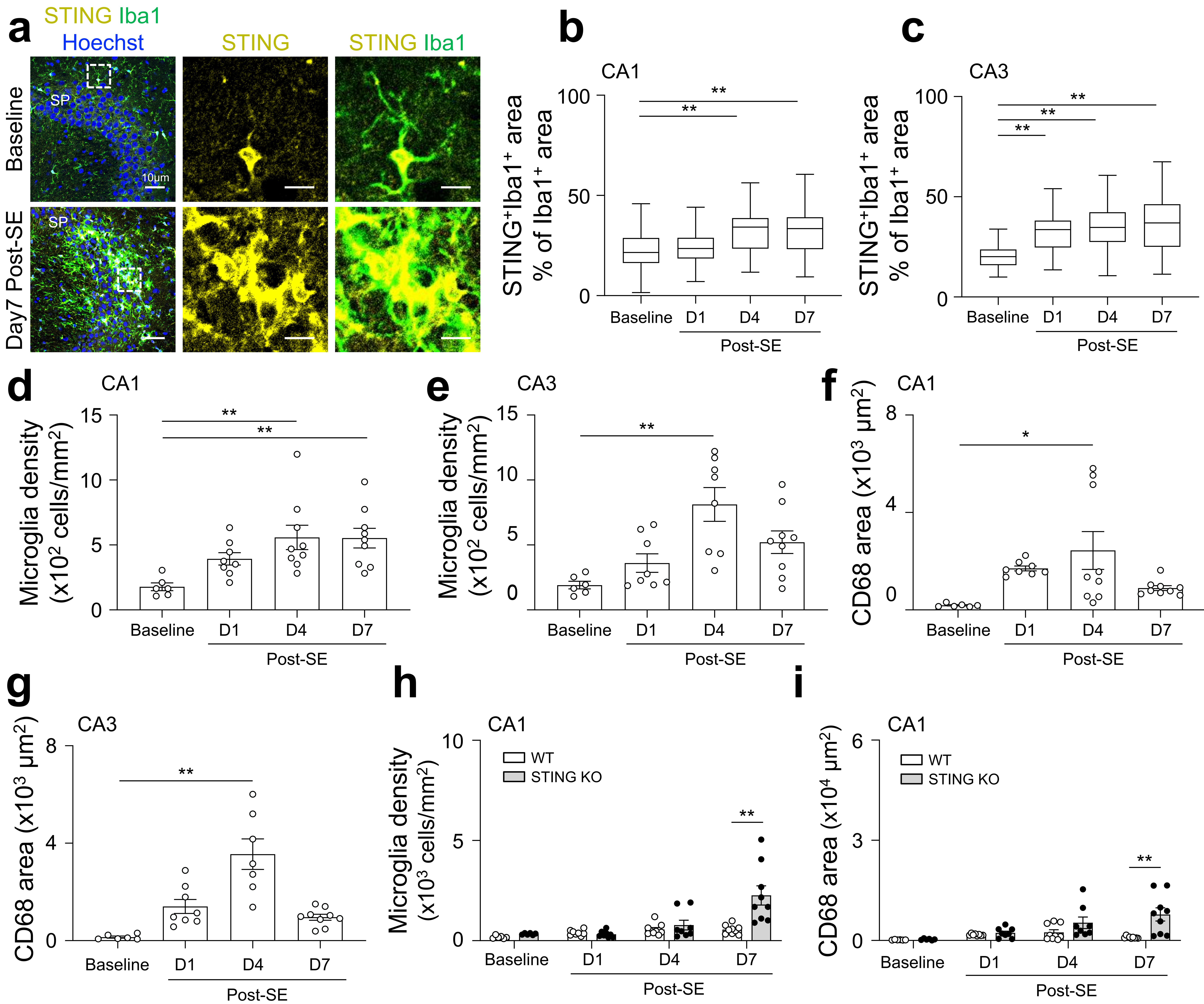


**Extended Data Fig. 1 Microglial and neuronal densities in the hippocampal CA regions of WT and STING-KO mice**

**a**, Representative images of the CA3 region of the hippocampus from WT mice under naïve baseline conditions and at 7 days post-SE. Areas enclosed by dashed rectangles are shown at higher magnification (right). Iba1 (green), STING (yellow), and NeuN (blue).

**b**, **c**, Percentage of $STING^{+}Iba1^{+}$ area relative to the total $Iba1^{+}$ area in individual microglia in the CA1 (b) and CA3 (c) regions.

**d, e**, Microglial density in the CA1 (d) and CA3 (e) regions in WT mice.

**f**, **g**, Quantification of CD68-positive area in the CA1 (f) and CA3 (g) regions in WT mice.

**h**, **i**, Microglial density (h) and CD68-positive area (i) in the CA1 region of WT and STING-KO mice.

Data in e and g are reproduced from Fig. 1f, h. Data are presented as mean ± s.e.m. Statistical analysis was performed using one-way ANOVA with Tukey's multiple-comparison test (b–g) or two-way ANOVA followed by Tukey's multiple-comparison test (h, i). $^{**}P < 0.01$ and $^{*}P < 0.05$. $n$ = 41–53 cells (b) and 38–51 cells (c) from 3 mice, and 6–9 regions from 3 mice per group (d–i).

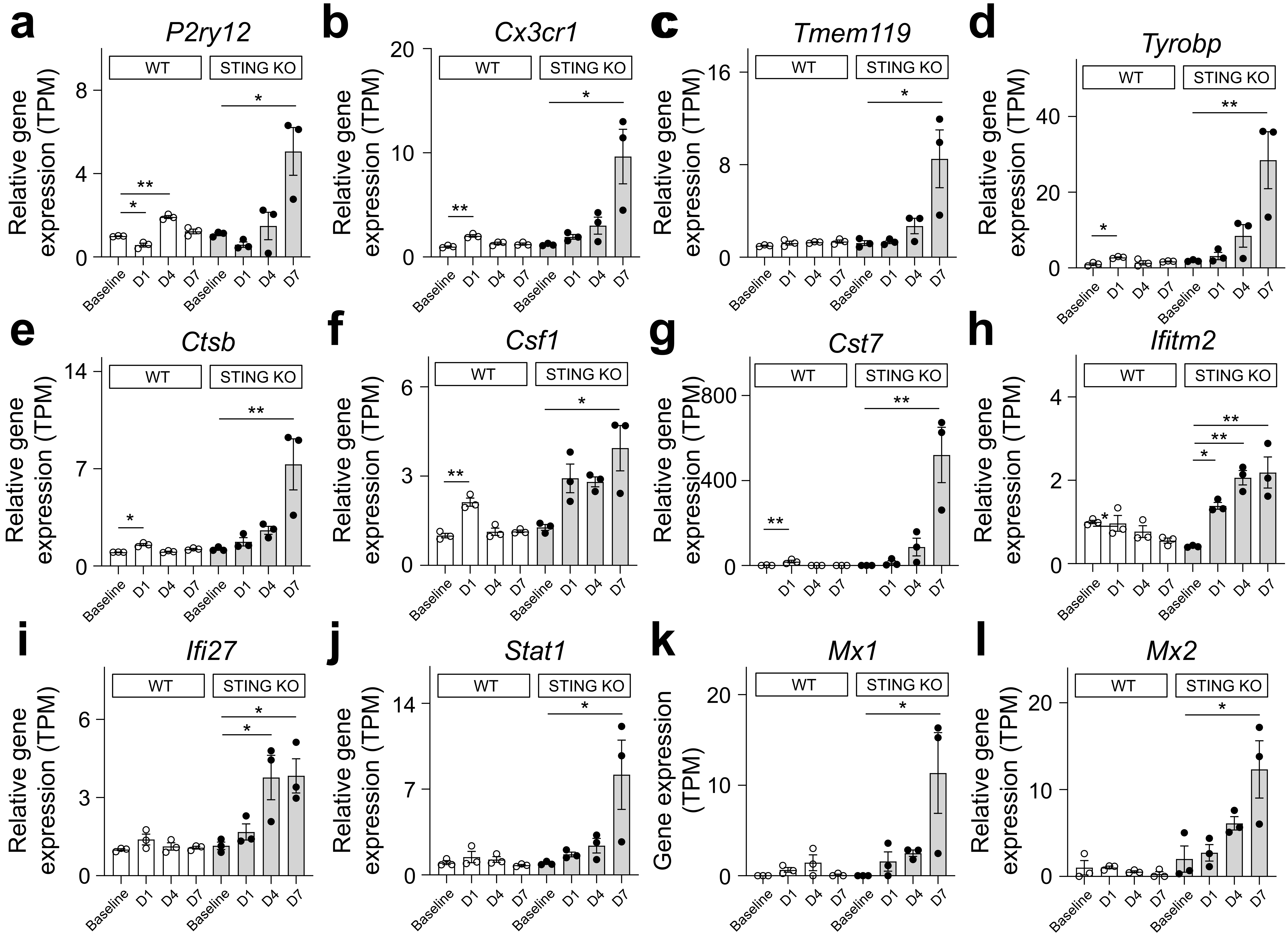


**Extended Data Fig. 2 Gene expression profiles in WT and STING-KO mice.**

mRNA expression of microglial homeostatic markers (a–c), activation-associated genes (d–g), and ISGs (h–l) in the hippocampus of WT and STING-KO mice. Panels show expression levels under naïve conditions and after SE within each genotype. Data are derived from Fig. 1a–d. Data are presented as mean ± s.e.m. Statistical analysis was performed using one-way ANOVA with Tukey's multiple-comparison test for within-genotype comparisons.

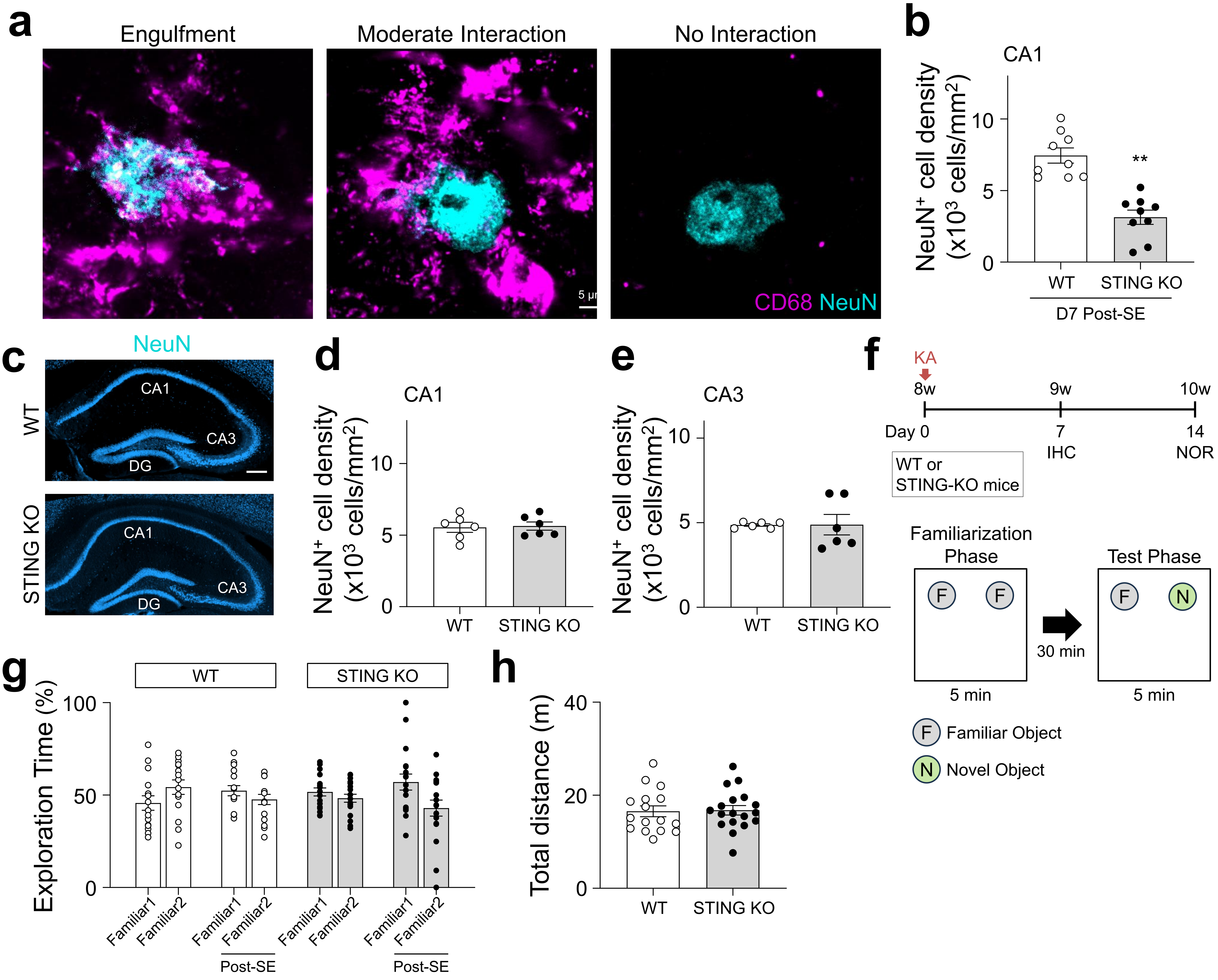


**Extended Data Fig. 3 NeuN⁺ cell densities and behavioral analyses in WT and STING-KO mice.**

**a**, Representative images illustrating phagocytic-stage classification based on the extent of overlap between CD68 and NeuN signals: Stage 1, no overlap (0%); Stage 2, minimal interaction (<10% overlap); Stage 3, moderate interaction (10–80% overlap); and Stage 4, engulfment (>80% overlap).

**b**, $NeuN^+$ cell density in the CA1 region at 7 days post-SE.

**c-e**, $NeuN^+$ cell densities in the CA1 and CA3 regions of WT and STING-KO mice under naïve conditions.

**f**, Experimental timeline and schematic of the NOR task.

**g**, Percentage of time spent exploring two familiar objects during the familiarization phase in WT and STING-KO mice.

**h**, Total distance traveled by WT and STING-KO mice during the test phase at 2 weeks post-SE.

Data are presented as mean ± s.e.m. Statistical analysis was performed using an unpaired Student's t-test (b, c–e, g) or two-way ANOVA followed by Tukey's multiple-comparison test (f). **P < 0.01 and *P < 0.05. *n* = 6–9 regions from 3 mice (b), 6 regions from 3 mice (c, d), and 16–18 mice (f, g).

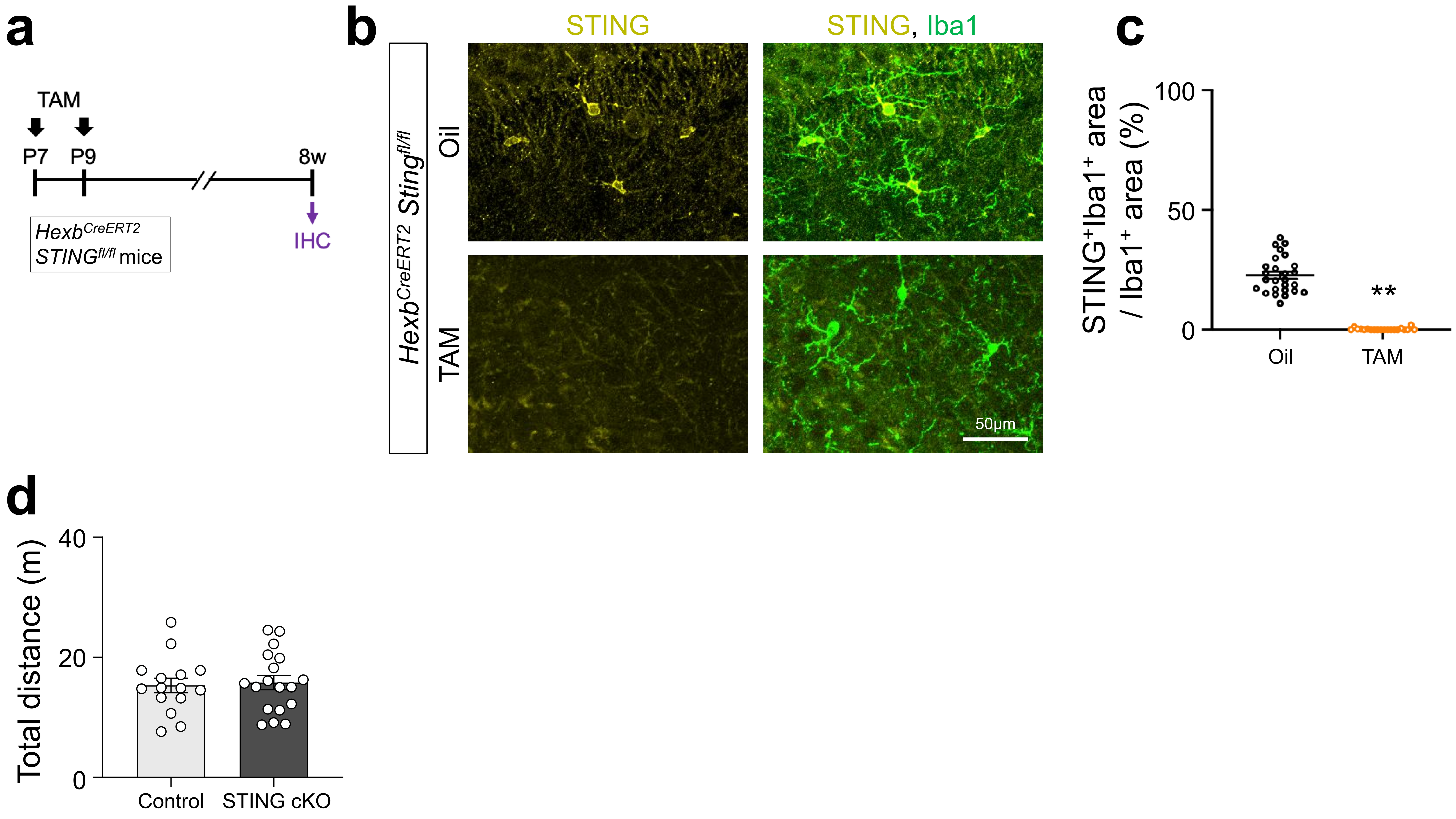


**Extended Data Fig. 4 Validation of conditional STING deletion in *Hexb$^{CreERT2}$ STING$^{fl/fl}$* mice**

**a**, Experimental timeline for tamoxifen (TAM) administration. *Hexb$^{CreERT2}$ STING$^{fl/fl}$* mice received TAM or vehicle (corn oil) at P7 and P9.

**b**, Representative images of the CA3 region of the hippocampus from *Hexb$^{CreERT2}$ STING$^{fl/fl}$* mice receiving TAM or vehicle. Iba1 (green) and STING (yellow).

**c**, Percentage of STING$^+$Iba1$^+$ area relative to the total Iba1$^+$ area in individual microglia.

**d**, Total distance traveled by control and STING cKO mice during the NOR test phase at 2 weeks post-SE.

Data are presented as mean ± s.e.m. Statistical analysis was performed using an unpaired Student's t-test. **P < 0.01. *n* = 20–26 microglia (c) and 14–18 mice (d).

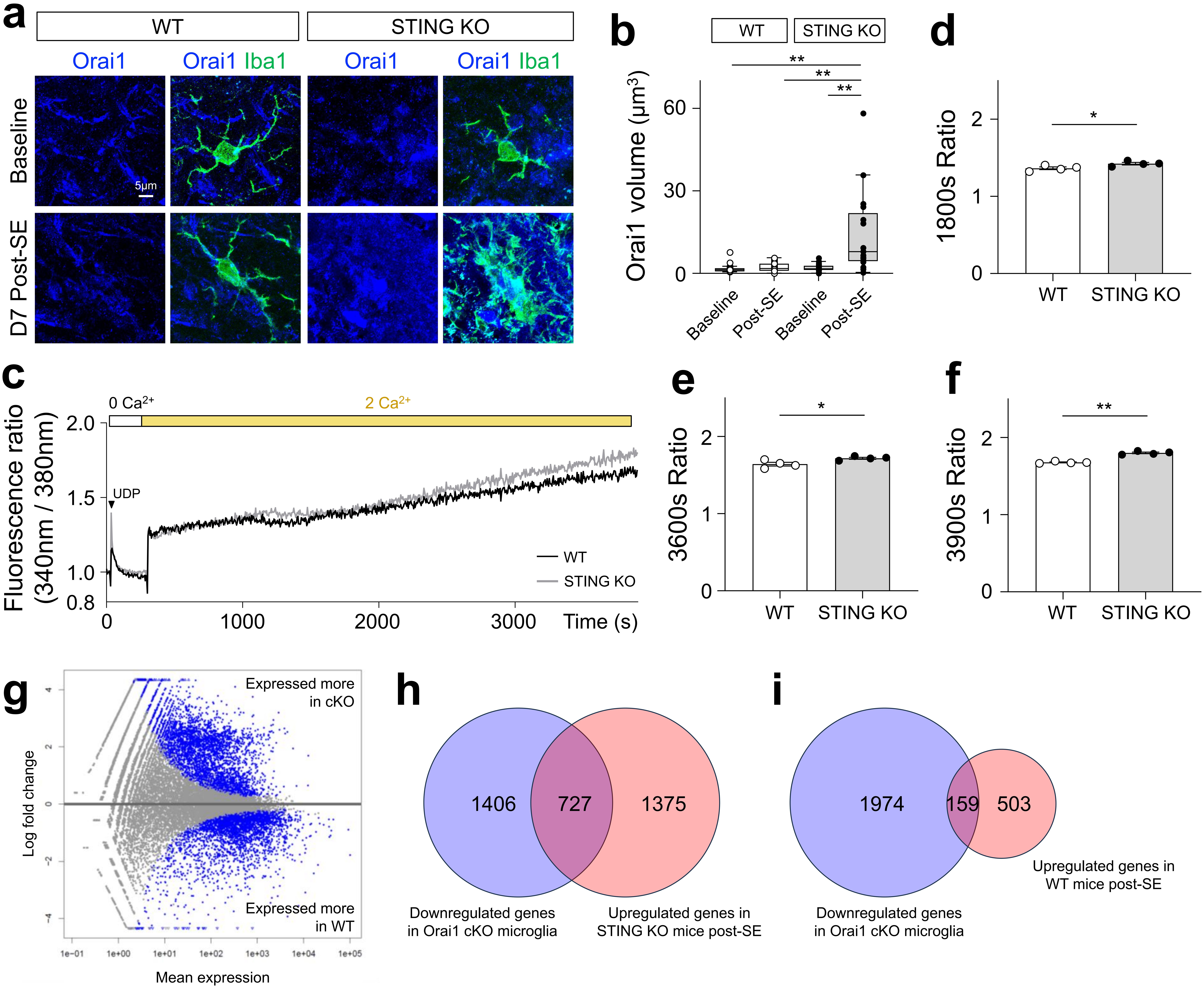


**Extended Data Fig. 5 Orai1 expression and SOCE activity in WT and STING-KO mice.**

**a**, Representative images of microglia in WT and STING KO mice under naïve conditions and at 7 days post-SE. Iba1 (green) and Orai1 (blue).

**b**, Quantification of Orai1-positive signal volume in individual microglia.

**c**, Time course of UDP-evoked SOCE in WT and STING-KO primary cultured microglia.

**d–f**, Fluorescence ratio at 1800 s (d), 3600 s (e), and 3900 s (f) after recording onset.

**g**, MA plot (log-fold change versus average expression) showing differential gene expression between WT and Orai1 cKO microglia after stimulation. Blue dots indicate DEGs (genes expressed at significantly higher or lower levels in WT compared with Orai1 cKO microglia), whereas gray dots indicate genes without significant differential expression between genotypes.

**h**, Venn diagram showing the overlap between genes downregulated in Orai1 cKO microglia and genes upregulated in STING-KO mice at 7 days post-SE compared with naïve controls. A total of 727 genes were shared.

**i**, Venn diagram showing the overlap between genes downregulated in Orai1 cKO microglia and genes upregulated in WT mice at 1 day post-SE compared with naïve controls. A total of 159 genes were shared.

For box plots, center lines indicate medians, box limits indicate the first and third quartiles, and whiskers extend to 1.5 × the interquartile range, as defined by Tukey. Statistical analysis was performed using one-way ANOVA with Tukey's multiple-comparison test (b) or an unpaired Student's t-test (d–f). **P < 0.01 and *P < 0.05. *n* = 17–20 microglia from 3 mice (b) and 4 independent cultures (c–f).

**a**

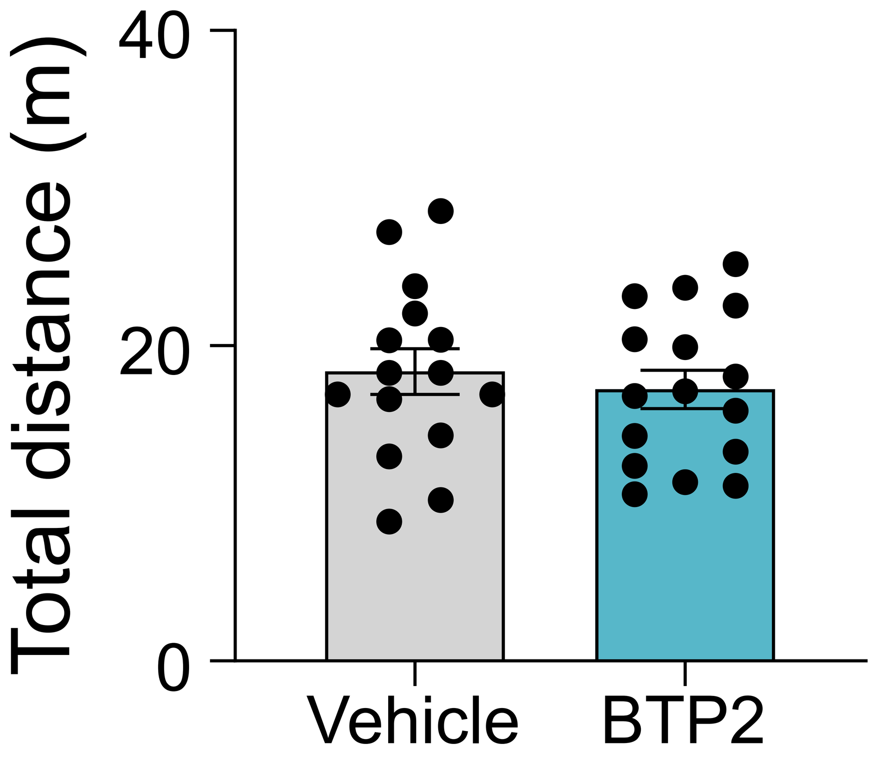


**Extended Data Fig. 6 Effects of BTP2 on locomotor activity in STING-KO mice.**

**a**, Total distance traveled by vehicle- or BTP2-treated STING-KO mice during the NOR test phase at 2 weeks post-SE.

Data are presented as mean ± s.e.m. Statistical analysis was performed using an unpaired Student's t-test. *n* = 15–16 mice.
